# Surface adhesion-dependent DNA extraction bias points to an “anvil effect” during mechanical lysis

**DOI:** 10.64898/2026.11.10.765706

**Authors:** Alex Batista Trentin, Roland C. Wilhelm

## Abstract

DNA extraction bias is usually treated as an intrinsic property of a cell’s envelope, size, or physiology rather than a feature of its physical location. Most soil bacteria live attached to surfaces, and attached cells should lyse preferentially during bead-beating if a bead striking a particle crushes the cell against it, as on an anvil, whereas a free cell must be caught between colliding beads. We tested this ‘anvil effect’ in sterile synthetic soil microcosms using isogenic strain pairs that differ in adhesion in two bacteria with different modes of attachment. *Agrobacterium tumefaciens* anchors with a unipolar polysaccharide and aggregates in an exopolysaccharide matrix, whereas *Caulobacter vibrioides* attaches individually by a polar holdfast. Within each strain we calculated the ratio of DNA yield from mechanical to chemical lysis, then compared adherent with non-adherent strains. In support of the anvil effect, mechanical lysis favored adherent cells in *A. tumefaciens* (2.0-fold) and *C. vibrioides* (2.6-fold). The route by which soil was transferred to the lysis tube altered the extent of bias observed, further indicating the importance of considering physical context during DNA extraction. Our synthetic soil was standardized and aggregate-free, leaving cells exposed on particle surfaces, making these results a direct test of the anvil effect. In natural soils, the aggregation, texture, and moisture may offset or amplify this effect. Surface attachment therefore biases soil DNA extraction, complicating corrections that treat extraction efficiency as a fixed, strain-specific coefficient.

**Importance:** Much of what is known about soil microbial communities comes from DNA extracted from soil, yet every extraction method has trade-offs in how well it breaks apart different kinds of cells. Most soil bacteria live attached to mineral particles, and we show that attachment itself changes how readily a cell releases its DNA. Compared with a chemical method, bead beating, a standard way of breaking cells open, recovered proportionally more DNA from strains that attach to particles than from matched strains that cannot. How soil was handled before extraction also changed the size of this bias. Extraction bias therefore depends on where a cell is, not only on what it is made of. Communities whose members shift between attached and free-living states could appear to change in composition when they have not, so these findings call for cautious interpretation, especially for populations known to adhere to surfaces.

## Background

DNA extraction is a key methodological decision point in any microbiome study (1–3), with choices affecting downstream results and ecological interpretation (4). Differences in extraction efficiency among diverse microbial envelopes and physiologies inherently bias the estimation of community composition, diversity, and differential abundance (5), with known trade-offs in the lysis of monoderm versus diderm populations (6) and cells of varying size (7). An estimated 80% of all terrestrial bacteria and archaea reside in surficial biofilms (8), yet whether surface attachment itself alters extraction efficiency has, to our knowledge, not been tested directly. Surface adhesion could bias extraction through several mechanisms (**Table S1**), but the most direct stems from particle collisions, where an attached cell is struck whenever a bead hits its particle, like an anvil, whereas a planktonic cell must be caught between colliding beads to lyse (9). Attached cells should therefore lyse preferentially during bead beating and be overrepresented in the resulting community profiles, unless their adhesive matrix resists disruption, as biofilms often do (10).

Soil is a prime context to test this ‘anvil effect,’ because the choice of extraction method strongly shapes soil community profiles (11, 12), and some of this variation may originate from the fate of surface-adherent soil populations during lysis. Here, we asked whether surface adhesion biases DNA recovery from a synthetic soil without aggregates, expecting that adherent cells would lyse more readily by mechanical than by chemical treatment relative to otherwise identical non-adherent cells. We compared isogenic strain pairs that differ in adhesion phenotype in two bacteria with contrasting modes of adhesion (**Fig. 1AB**), namely *Agrobacterium tumefaciens*, which anchors by a unipolar polysaccharide (UPP; wild type versus a UPP-deficient mutant) and embeds in a bulk exopolysaccharide matrix (13, 14), and *Caulobacter vibrioides*, which attaches by a discrete polar holdfast (holdfast-overproducing Δ*hfiA* versus holdfast-deficient Δ*hfsDAB*) and forms surface monolayers and rosettes rather than a thick matrix (15, 16). Normalizing mechanical to chemical DNA yield within each strain to account for differences in cell load, we compared phenotypes by dividing the adherent strain’s ratio by that of the non-adherent strain (ratio of ratios, ROR), where an ROR > 1 means that, relative to chemical lysis, mechanical lysis favors adherent cells (**Fig. 1C**). To test whether any bias reflected adhesion rather than sample handling, we repeated the comparison under different handling methods.

**Figure 1.**
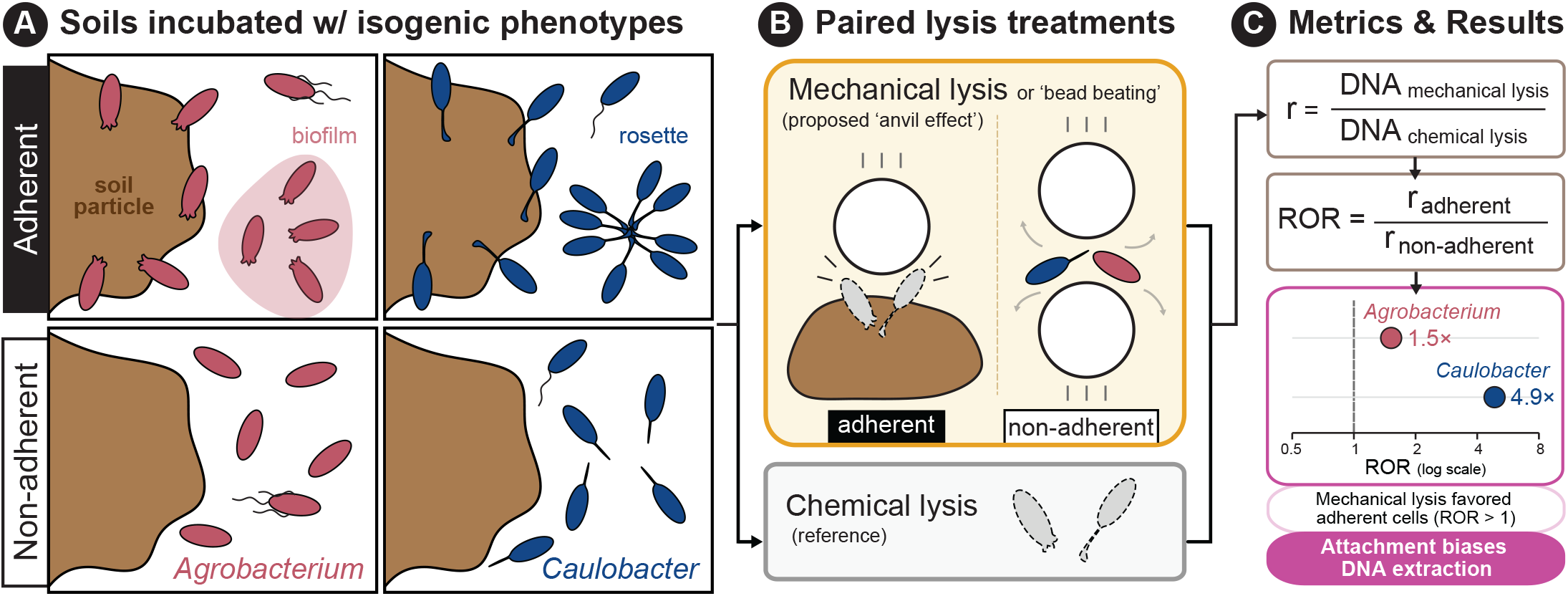
An overview of the experimental design testing the proposed anvil effect. (A) Sterile synthetic soil microcosms were inoculated with isogenic adherent and non-adherent strains of *Agrobacterium tumefaciens* and *Caulobacter vibrioides*. (B) Soil from each microcosm was split between mechanical lysis by bead beating and chemical lysis as the reference. (C) The ratio of mechanical to chemical DNA yield (r) was calculated for each microcosm, and the ratio of ratios (ROR) was used to compare r between adherent and non-adherent strains. (Alt text). A three-panel schematic. In the first panel, soil particles carry attached *Agrobacterium* and *Caulobacter* cells, while non-adherent strains float free. The second panel shows mechanical lysis by bead beating, in which a bead crushes an attached cell against its particle (the proposed anvil effect) while free cells are swept aside, alongside chemical lysis as the reference. The third panel shows how r is calculated as the ratio of DNA yield from mechanical to chemical lysis, and how the ratio of ratios (ROR) divides r for the adherent strain by r for its non-adherent isogenic partner. A plot of the resulting RORs shows values above 1 for both Agrobacterium (1.5) and Caulobacter (4.9).

### Strains, microcosms, and DNA extractions

Experimental DNA extractions were performed on cultures of *A. tumefaciens* C58 (wild type, adherent) and its Δ*upp* mutant PMM26 (non-adherent), and *C. vibrioides* (syn. *C. crescentus*) CB15 Δ*hfiA* (FC1365, adherent) and Δ*hfsDAB* (YB2857, non-adherent). Growth media, growth rates, and microscope images available in **Supplementary Methods** (**Figs. S1** and **S2**). Microcosms with 3 g of synthetic soil (70% quartz silt, 20% sand, and 10% muscovite clay), sterilized by triple autoclaving (17), were inoculated with 5 mL of washed cells (OD^600^ = 0.1) and shaken at 200 rpm for 24 h at room temperature. Uninoculated controls carried through every step yielded no quantifiable DNA. Soil from each microcosm was split between mechanical lysis by bead beating (6.0 m s^−1^, 40 s) and chemical lysis with Triton X-100, SDS, and proteinase K at 56°C (18, 19), after which both lysates were purified identically with the DNeasy PowerSoil Pro kit (Qiagen) and quantified with the Qubit dsDNA HS assay. Because washing removed most non-adherent cells, only soils incubated with adherent strains were washed. In Variation 1 (n = 12 microcosms per strain), adherent soil was pelleted, washed, and weighed (250 mg wet soil) into the lysis tube, whereas the non-adherent slurry was vortexed and pipetted (350 µL). Variation 2 (n = 4 microcosms per strain per route) compared two routes for transferring soil from the same microcosms, either pelleted and weighed (250 mg; weighed-soil route) or pipetted as 350 µL of vortexed suspension, the adherent soil having first been pelleted, washed, and resuspended in 5 mL of water (resuspended-soil route). For each microcosm, ‘r’ is the ratio of DNA yield from mechanical to chemical lysis. Within each bacterium, log r was compared between strains with Welch’s t-tests, and the back-transformed difference gave the ROR and its 95% CI. Changes in ROR between routes were tested as differences in log ROR (**Supplementary Methods**). Claude (Anthropic, CA) assisted with scripting and text editing.

### Results and Discussion

Relative to chemical lysis, mechanical lysis recovered proportionally more DNA from adherent than from non-adherent phenotypes in both bacteria, as predicted by the anvil effect. In Variation 1 (n = 12 per strain), the ROR exceeded 1 for both *A. tumefaciens* (ROR = 2.03; 95% CI, 1.38–2.98; P = 0.001) and *C. vibrioides* (ROR = 2.58; 95% CI, 1.87–3.55; P < 0.001; **Fig. 2A**). Because handling differed between phenotypes in Variation 1, Variation 2 applied the same handling to both phenotypes, comparing two routes for transferring an equivalent amount of soil from the same microcosms to the lysis tube (n = 4 per strain, per route). In the ‘weighed-soil route,’ in which soil was pelleted and the wet soil weighed into the lysis tube, the bias toward adherent cells persisted, with RORs of 1.52 for *A. tumefaciens* (95% CI, 1.14–2.03; P = 0.013) and 4.86 for *C. vibrioides* (95% CI, 2.14–11.03; P = 0.005; **Fig. 2B**). The ROR was higher for *C. vibrioides* than for *A. tumefaciens* in Variation 1 and in the weighed-soil route (**Fig. 2**). Surface attachment therefore favored DNA recovery by mechanical lysis, contrary to the suggestion that adsorption to soil particles protects cells from lysis (20), at least during bead beating, and the magnitude of this bias differed between the two modes of adhesion.

**Figure 2.**
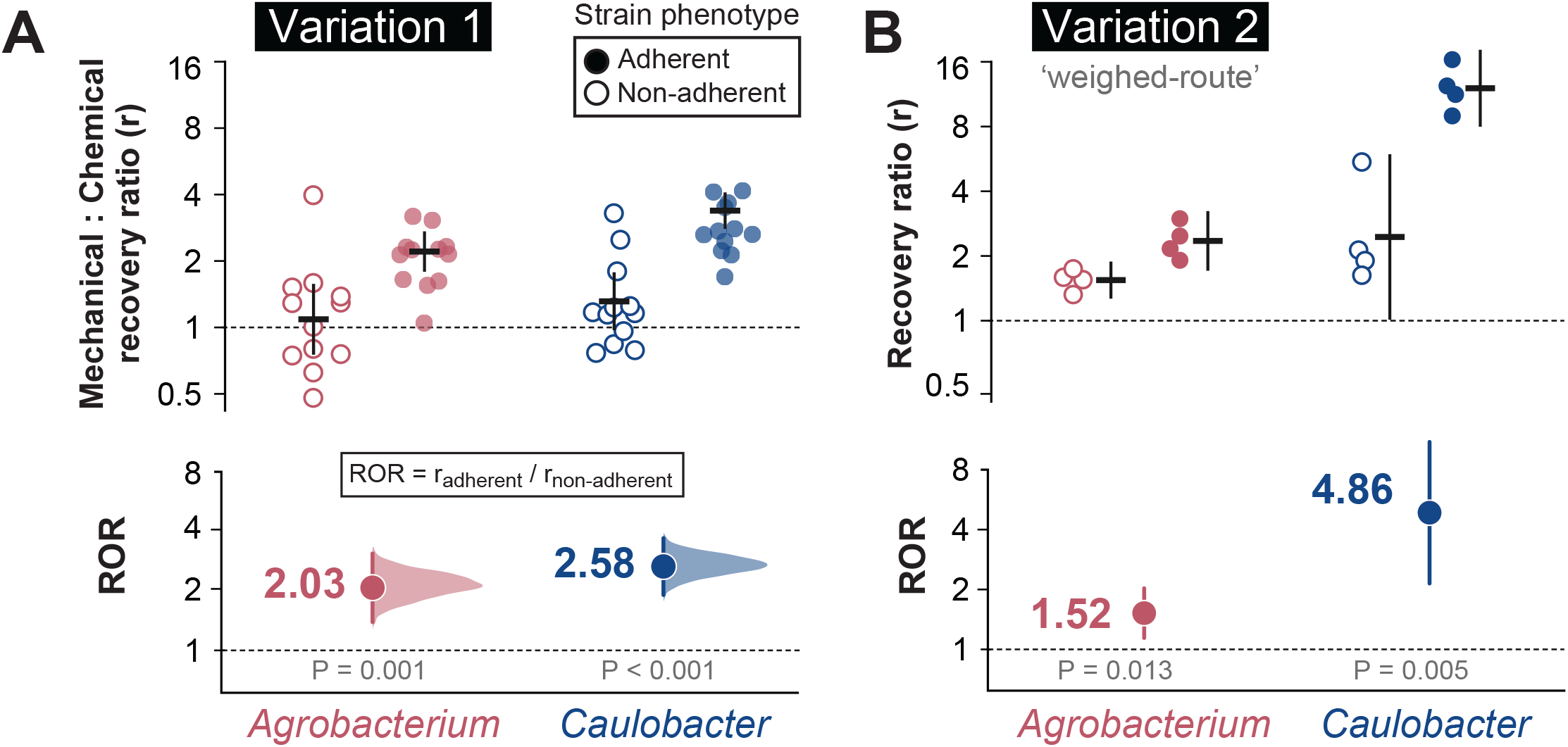
Mechanical lysis favors DNA recovery from adherent phenotypes of *Agrobacterium tumefaciens* and *Caulobacter vibrioides*. Upper panels show the ratio of DNA yield from mechanical to chemical lysis (r) for adherent (filled; *A. tumefaciens* wild type, *C. vibrioides* Δ*hfiA*) and non-adherent (open; *A. tumefaciens* Δ*upp, C. vibrioides* Δ*hfsDAB*) strains. Points correspond to soil from individual microcosms and crosses show geometric means ± 95% CI. Lower panels show the ratio of ratios, ROR = r_adherent_ / r_non-adherent_, ± 95% CI, where ROR > 1 indicates that mechanical lysis, relative to chemical lysis, favors adherent cells. In panel (A) half-violins show the bootstrap distribution of the ROR for Variation 1 (n = 12 microcosms per strain). In panel (B), the same visualizations using soils sampled according to Variation 2, in which soil from both phenotypes was pelleted and weighed wet into the lysis tube (‘weighed-soil route’; n = 4 microcosms per strain). Dashed lines mark r = 1 and ROR = 1. P values are from Welch’s t-tests on log-transformed r. Y-axes are displayed on a log2 scale, with tick labels shown in the original ratio units. (Alt text). Two-panel figure with dot plots of mechanical-to-chemical DNA recovery ratios for adherent and non-adherent *Agrobacterium* and *Caulobacter*, with lower summary plots showing ratios of ratios all above 1, equaling 2.03 and 2.58 in Variation 1 for Agrobacterium and Caulobacter, respectively and 1.52 and 4.86 in the weighed-soil route of Variation 2.

In the resuspended-soil route of Variation 2, the adherence bias persisted in *A. tumefaciens* (ROR = 3.14; 95% CI, 1.22-8.06; P = 0.029; **Fig. 3A**) but disappeared in *C. vibrioides*, whose ROR fell from 4.86 in the weighed-soil route to 1.01 after resuspension (95% CI, 0.76–1.35; P = 0.909; **Fig. 3A**). In adherent *C. vibrioides*, resuspension lowered DNA recovery by mechanical lysis 2.43-fold and raised recovery by chemical lysis 2.43-fold, whereas neither changed appreciably in the non-adherent strain (**Fig. 3B**), indicating that the effect of the transfer route was specific to the adherent strain. Improved reagent access is an unlikely cause, because soil was resuspended and vortexed during chemical lysis in both routes of Variation 2. More plausibly, the pipetted aliquot sampled a different mix of cells and particle sizes following resuspension. Adherent cells that co-sediment with the coarse particles would be under-represented, leaving fewer and smaller particles to act as anvils, and unattached cells would be over-represented in the resuspended material, for which chemical lysis is relatively more effective. The persistence of the adherence bias in *A. tumefaciens* may be a facet of its exopolysaccharide matrix (**Fig. S1**), with matrix-bound cells behaving as lightweight particles during resuspension, preserving some of the anvil effect as well as chemical shielding, which drive the ROR.

**Figure 3.**
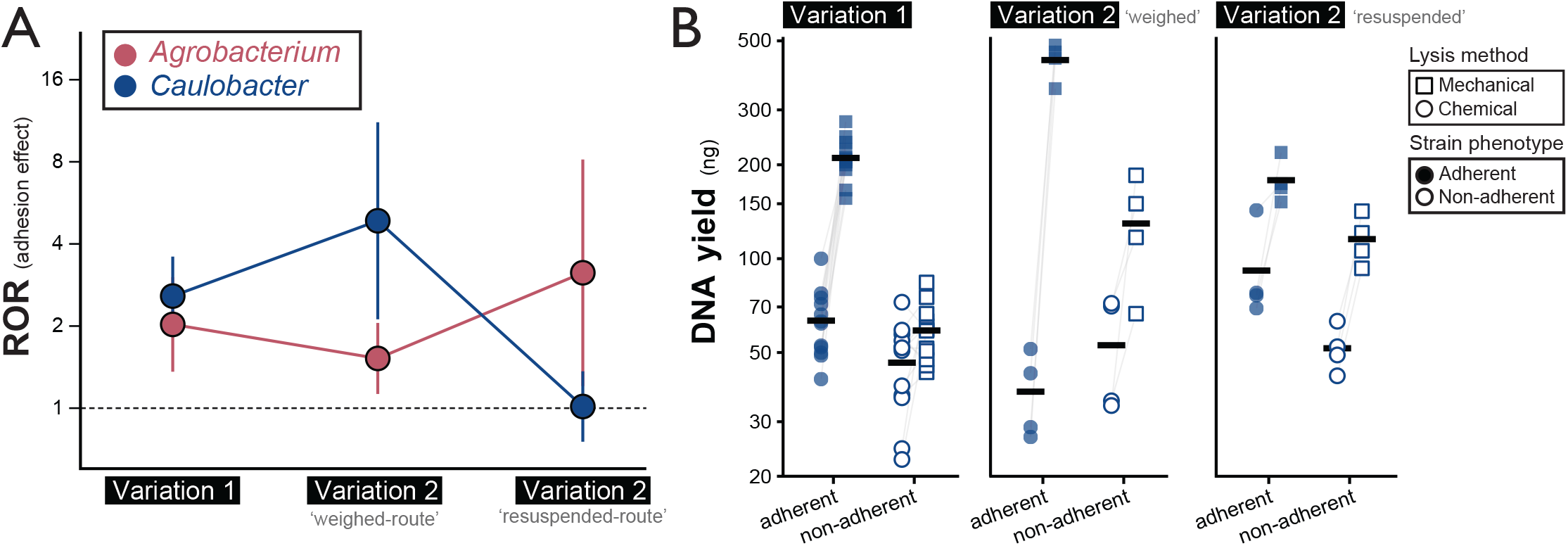
Resuspending soil before transferring to the lysis tube removes the adhesion bias in *Caulobacter vibrioides* but not in *Agrobacterium tumefaciens*. In panel (A), ROR for *A. tumefaciens* (red) and *C. vibrioides* (blue) under Variation 1 and the two routes of Variation 2. These routes transferred soil from the same microcosms to the lysis tube either as pelleted, weighed wet soil (weighed-soil route) or as a pipetted aliquot of soil suspension (resuspended-soil route). The dashed line marks ROR = 1 (no adhesion effect). In panel (B), DNA yields from adherent (filled; Δ*hfiA*) and non-adherent (open; Δ*hfsDAB*) *C. vibrioides* after chemical (circles) or mechanical (squares) lysis. Points are individual extractions, gray lines join the paired extractions from each microcosm, and black bars show geometric means. n = 12 (Variation 1) or 4 (each Variation 2 route) microcosms per strain. Y-axes are logarithmically scaled; panel A is displayed on a log2 scale and panel B on a log10 scale, with tick labels shown in the original units. (Alt text). Two-panel figure. Adhesion-effect ratios for *Agrobacterium* stay above 1 across three handling conditions (2.03, 1.52, 3.14), whereas *Caulobacter*’s fall from 2.58 and 4.86 to 1.01 after resuspension. Paired *Caulobacter* DNA yields show resuspension raised chemical and lowered mechanical recovery mainly in the adherent strain.

These results suggest that surface-adherent populations are likely overrepresented in soil DNA extracted by mechanical lysis, to a degree that depends on adhesion architecture and on the physical context of the sample in the lysis tube. Our synthetic soil had a silt loam texture but no aggregates, making it a purer test of the anvil effect. Cells could attach to particle surfaces but could not shelter inside aggregates and avoid impact (20). In natural soils, such shelter, and the other mechanisms in **Table S1**, may offset or enhance the anvil effect. Because adhesion-dependent bias is also likely to vary with texture, aggregation, moisture and the mode of particle association, no single corrective measure will suffice. Thus, calibration strategies that treat extraction efficiency as a fixed, taxon-specific coefficient may fail in soil (21), since a correction factor estimated from freely suspended cells, or from a particular soil environment, may not transfer to other contexts. Mock communities used to quantify extraction biases (22) could help address this by including isogenic adherent and non-adherent strain pairs, alongside members chosen for cell envelope or 16S rRNA gene copy number. Likewise, because a single pre-lysis step altered the bias substantively, the composition of samples transferred to the lysis tube (wet soil, dry soil etc.) should be standardized within an experiment and reported.

## Acknowledgements

We thank Yves Brun and Gail Hardy for providing the *Caulobacter vibrioides* Δ*hfsDAB* mutant, Aretha Fiebig for providing the Δ*hfiA* mutant, and Clay Fuqua for providing the *Agrobacterium tumefaciens* wild-type strain and *Δupp* mutant. We thank Kaitlyn Annunziata for characterizing the growth rates and morphologies of *Caulobacter* and *Agrobacterium*. Funding support for ABT was provided by the Brazilian Ministry of Education and the Coordenação de Aperfeiçoamento de Pessoal de Nível Superior (CAPES) Fellowship Program – Finance Code 001 (https://ror.org/00x0ma614). Partial funding support for RCW was provided by the USDA National Institute of Food and Agriculture (NIFA; https://ror.org/05qx3fv49) Hatch Grant program (IND90024429), administered by Purdue University (https://ror.org/05p8z3f47).

## Data Availability

The DNA yield data, metadata, and the R code for all analyses and figures are available in the Open Science Framework repository at https://doi.org/10.17605/OSF.IO/A7EMH.

## References

1. Sui H, Weil AA, Nuwagira E, Qadri F, Ryan ET, Mezzari MP, Phipatanakul W, Lai PS. 2020. Impact of DNA Extraction Method on Variation in Human and Built Environment Microbial Community and Functional Profiles Assessed by Shotgun Metagenomics Sequencing. Front Microbiol 11:953.

2. Browne DJ. 2026. The Great Microbiome Extraction Anomaly: How Extraction Bias Distorts Microbial Community Profiles. Environmental DNA 8:e70276.

3. Brooks JP, Edwards DJ, Harwich MD, Rivera MC, Fettweis JM, Serrano MG, Reris RA, Sheth NU, Huang B, Girerd P, Strauss JF, Jefferson KK, Buck GA. 2015. The truth about metagenomics: quantifying and counteracting bias in 16S rRNA studies. BMC Microbiol 15:66.

4. Reese J-O, Castro Lund IM, Haugen HJ, Saragliadis A, Lyngstadaas SP, Linke D. 2026. The impact of DNA extraction methods on species quantification and apparent community composition of in vitro oral biofilms. FEMS Microbiology Letters 373:fnag066.

5. Dopheide A, Xie D, Buckley TR, Drummond AJ, Newcomb RD. 2019. Impacts of DNA extraction and PCR on DNA metabarcoding estimates of soil biodiversity. Methods Ecol Evol 10:120–133.

6. Rauer L, De Tomassi A, Müller CL, Hülpüsch C, Traidl-Hoffmann C, Reiger M, Neumann AU. 2025. De-biasing microbiome sequencing data: bacterial morphology-based correction of extraction bias and correlates of chimera formation. Microbiome 13:38.

7. Moré MI, Herrick JB, Silva MC, Ghiorse WC, Madsen EL. 1994. Quantitative cell lysis of indigenous microorganisms and rapid extraction of microbial DNA from sediment. Appl Environ Microbiol 60:1572–1580.

8. Flemming H-C, Wuertz S. 2019. Bacteria and archaea on Earth and their abundance in biofilms. Nat Rev Microbiol 17:247–260.

9. Melendres A, Honda H, Shiragami N, Unno H. 1991. A kinetic analysis of cell disruption by bead mill. The influence of bead loading, bead size and agitator speed. Bioseparation 2:231–6.

10. Peterson BW, He Y, Ren Y, Zerdoum A, Libera MR, Sharma PK, Van Winkelhoff A-J, Neut D, Stoodley P, Van Der Mei HC, Busscher HJ. 2015. Viscoelasticity of biofilms and their recalcitrance to mechanical and chemical challenges. FEMS Microbiology Reviews 39:234–245.

11. Cardoni M, Nguyen H, Royle J, Bennett JA. 2026. Lost in Extraction: Uncovering Bias in Soil Microbial Community Profiles. Applied and Environmental Soil Science 2026:5563719.

12. Zielińska S, Radkowski P, Blendowska A, Ludwig-Gałęzowska A, Łoś JM, Łoś M. 2017. The choice of the DNA extraction method may influence the outcome of the soil microbial community structure analysis. MicrobiologyOpen 6:e00453.

13. Xu J, Kim J, Koestler BJ, Choi J, Waters CM, Fuqua C. 2013. Genetic analysis of A grobacterium tumefaciens unipolar polysaccharide production reveals complex integrated control of the motile-to-sessile switch. Molecular Microbiology 89:929–948.

14. Heindl JE, Wang Y, Heckel BC, Mohari B, Feirer N, Fuqua C. 2014. Mechanisms and regulation of surface interactions and biofilm formation in Agrobacterium. Front Plant Sci 5.

15. Hershey DM, Porfírio S, Black I, Jaehrig B, Heiss C, Azadi P, Fiebig A, Crosson S. 2019. Composition of the Holdfast Polysaccharide from Caulobacter crescentus. J Bacteriol 201.

16. Entcheva-Dimitrov P, Spormann AM. 2004. Dynamics and Control of Biofilms of the Oligotrophic Bacterium Caulobacter crescentus. J Bacteriol 186:8254–8266.

17. King WL, Grandinette EM, Trase O, Rolon ML, Salis HM, Wood H, Bell TH. 2024. Autoclaving is at least as effective as gamma irradiation for biotic clearing and intentional microbial recolonization of soil. mSphere 9:e00476–24.

18. Zhou J, Bruns MA, Tiedje JM. 1996. DNA recovery from soils of diverse composition. Applied and Environmental Microbiology 62:316–322.

19. Lever MA, Torti A, Eickenbusch P, Michaud AB, Šantl-Temkiv T, Jørgensen BB. 2015. A modular method for the extraction of DNA and RNA, and the separation of DNA pools from diverse environmental sample types. Front Microbiol 6:476.

20. Frostegård Å, Courtois S, Ramisse V, Clerc S, Bernillon D, Le Gall F, Jeannin P, Nesme X, Simonet P. 1999. Quantification of Bias Related to the Extraction of DNA Directly from Soils. Appl Environ Microbiol 65:5409–5420.

21. McLaren MR, Willis AD, Callahan BJ. 2019. Consistent and correctable bias in metagenomic sequencing experiments. eLife 8:e46923.

22. Bokulich NA, Rideout JR, Mercurio WG, Shiffer A, Wolfe B, Maurice CF, Dutton RJ, Turnbaugh PJ, Knight R, Caporaso JG. 2016. mockrobiota: a Public Resource for Microbiome Bioinformatics Benchmarking. mSystems 1:e00062–16.

